# Identification and characterization of a non-peptidic cyclophilin ligand with antiviral activity against feline and porcine α-coronaviruses

**DOI:** 10.1101/2025.05.24.655938

**Authors:** Manon Delaplace, Mantasha Khan, Hélène Huet, Laurent Softic, Lionel Bigault, Jean-François Guichou, Abdelhakim Ahmed-Belkacem, Quentin Nevers, Sophie Le Poder

## Abstract

Coronaviruses (CoVs) are emerging pathogens that have been extensively studied over the last 20 years and can cause acute respiratory diseases in humans, as exemplified by the SARS-CoV-2 pandemic. Coronaviruses are also known for their importance in veterinary medicine, being responsible for severe pathologies in pets and livestock. These include Feline Infectious Peritonitis Virus (FIPV), which causes a fatal disease in cats. In livestock, porcine coronaviruses such as Transmissible Gastroenteritis Virus (TGEV) and Porcine Epidemic Diarrhoea Virus (PEDV) are the causative agents of an acute enteric disease in piglets with a high mortality rate and a significant impact on the pork industry. In addition, animal coronaviruses may represent a zoonotic reservoir. Therefore, efficient antiviral strategies are required to inhibit the multiplication of coronaviruses infecting various animal species. Here, we synthesized 20 small-molecule ligands that target cyclophilins, a family of cellular chaperons hijacked by several viruses including CoVs. We screened their antiviral activity against feline and porcine alpha-CoVs, and identified a compound, F83233, as a potent inhibitor of FIPV, TGEV and PEDV replication at micromolar concentrations that was effective in feline, porcine and simian cells. As cyclophilins are highly conserved among mammals, F83233 could be a promising antiviral to treat different animal and zoonotic coronaviruses.

## Introduction

Coronaviruses (CoVs) are enveloped viruses with a single-stranded RNA genome of positive polarity. Inside the *Coronaviridae* family, the sub-family *Orthocoronavirinae* is composed of four genera: *Alpha (α)-, Beta (β)-, Gamma (γ)*- and *Delta (δ)-coronavirus*. α-CoVs and β-CoVs mainly infect mammals while γ-CoVs and *δ*-CoVs mostly infect birds with a few exceptions of mammals^1^. CoVs have a huge impact on public health: since 2002, they have been recognized as emerging and important pathogens in humans. Indeed, three zoonotic CoVs belonging to the β-CoV genus inducing severe pulmonary diseases have emerged in humans: SARS-CoV in 2002, MERS-CoV in 2012 and SARS-CoV-2 in 2019^2–4^. The emergence of these viruses, especially SARS-CoV-2, the etiological agent of the COVID-19, has led to a fantastic increase in knowledge about β-CoVs biology and pathophysiology.

In veterinary medicine also, CoVs are well known to trigger the development of various diseases, exhibiting sometimes a complex pathophysiology with a tropism which is not restricted to the respiratory tract^5,6^. The α-CoV genus contains an important number of pathogens that infect domestic and livestock animals, and effective treatments or vaccines are often missing for these viruses. Among them, FIPV (Feline Infectious Peritonitis Virus) causes a systematically fatal disease in cats, feline infectious peritonitis (FIP)^7^. FIP is considered one of the leading causes of death in communal cat groups with a 100% lethality rate, although treatment with GS-441524 now constitutes an excellent therapeutic strategy^8,9^. Feline CoVs (FCoV) are classified into two biotypes: Feline enteric CoV (FeCV) and Feline Infectious Peritonitis Virus (FIPV). FeCV is endemic in cats and avirulent, inducing mild or subclinical digestive symptoms. FIP disease is caused by virulent FIPV strains^7^. While FeCV has a strict intestinal tropism, FIPV is able to infect monocytes and macrophages^10^, allowing the virus to spread in various organs leading to a systemic infection. Beyond biotypes, FCoV is also subdivided, based on serological responses, into two serotypes: FCoV-I, the source of most natural infection amongst cats^11^, and FCoV-II resulting from recombination between FCoV-I and the canine coronavirus CCoV-II^12^. In contrast to FCoV-I, FCoV-II replicates readily in feline cell lines such as CRFK cells (Crandell-Reed Feline Kidney).

Several α-CoVs are also threatening farms and livestock animals. For example, six CoVs can infect pigs, among them four induce a clinically undistinguishable severe digestive disease, with case-fatality ratios of 80-100% in piglets that are less than 10-days^13,14^. Transmissible Gastroenteritis Virus (TGEV) is the prototype porcine CoV that shares an important sequence homology with FIPV. Interestingly, TGEV also exhibits a dual tropism *in vivo*, for intestinal and respiratory epithelia^5^. In the 1980’s, a deletion in the N-terminal domain of the Spike protein has led to the global emergence of a TGEV variant called PRCV (Porcine Respiratory Coronavirus) that lost the enteric tropism and virulence^5^. As antibodies elicited against PRCV protect against TGEV, the latter is now well controlled. Porcine Epidemic Diarrhoea Virus (PEDV) has emerged in the 1970’s: ancestral PEDV strains have been contained by vaccines, but PEDV is re-emerging since the 2010’s with the presence of new strains that render the vaccines less effective^15–17^. Re-emergence of PEDV in the United States in 2013 led to the loss of 10% of the pig herd with huge economic impact^18–20^. These PEDV strains now globally circulate^21^. Other porcine CoVs such as SADS-CoV (Swine Acute Diarrhoea Syndrome coronavirus) and PDCoV (porcine *Deltacoronavirus*) are emerging viruses with a documented zoonotic potential^22,23^. Considering the impressive capacity of CoVs to jump across the species barrier^24^ and the regular and severe resulting epidemics, it is of utmost importance to find, not only treatment for a specific pathogenic animal CoV, but also antiviral strategies that could be applied to a broad spectrum of CoVs, therefore preventing future emerging viruses.

All along the CoV life cycle^25^, viral proteins and viral RNAs are involved in interactions with host cellular factors that can facilitate or restrict the replication of CoVs^26– 30^. Among them are cyclophilins, conserved cellular proteins present in both prokaryotes and eukaryotes^31^. Cyclophilins share a common peptidyl-prolyl cis/trans isomerase domain (PPIase)^31^ which catalyzes the interconversion of proline configuration^32^. Cyclophilins are known to play critical roles in the replication of different viruses^33^, including DNA viruses^34^ as well as negative- and positive-sense RNA viruses^35,36^. They have also been shown to play a role in CoV replication, although their precise involvement along the life cycle is unclear^37^. Cyclosporin A (CsA), a macrocyclic inhibitor of cyclophilins, as well as its non-immunosuppressive derivatives such as alisporivir (ALV) inhibit CoV replication of different genera^38–40^. Gene knock-out or siRNA of cyclophilins render cells resistant to several CoV infections^41,42^. Cyclophilins have also been identified as interactors of several viral proteins of CoVs including SARS-CoV^43^ and HCoV-229E^44^.

We present here the characterization of the antiviral effect of small-molecule cyclophilin inhibitors^45^ (“ SMCypI” ) against a feline coronavirus of major veterinary interest, FIPV, and two enteric porcine CoVs, TGEV and PEDV. Following the synthesis and the screening of 20 SMCypI, we identified a compound referred as F83233 as a potent anti-FIPV compound in feline cells. This molecule also potently inhibited the multiplication of TGEV and PEDV in cells from pigs and monkeys, demonstrating its potential for the development of optimized antivirals effective to treat CoV diseases in animals, and to prevent the zoonotic risk caused by these pathogens.

## Materials and Methods

### Cells and virus strains

CRFK, ST, PK15 and Vero cells were cultivated at 37°C with 5% CO_2_ in Dulbecco’s modified Eagle’s medium (DMEM, Gibco) supplemented with 10% foetal bovine serum, 1% penicillin-streptomycin, 1% sodium pyruvate and 1% non-essential amino acids.

The serotype II FIPV 79-1146 was amplified on feline CRFK cells. Transmissible gastroenteritis virus (TGEV, Purdue strain) was amplified on porcine ST cells. Porcine epidemic diarrhoea virus (PEDV, CV777 strain) was amplified on simian Vero cells. Supernatants were harvested and ultracentrifugated. Infectious titers were determined using end-point dilution assays.

### PPIase enzyme assay

Human cyclophilin A was purified and its PPIase activity in absence or presence of SMCypI was measured as previously described^45^. Briefly, Cyclophilin A PPIase activity was measured at 20°C using the standard chymotrypsin-coupled assay. The assay buffer (25 mM Hepes and 100 mM NaCl, pH 7.8) and purified cyclophilin A were pre-cooled to 4°C. Then, 5 ml of 50 mg/ml chymotrypsin in 1 mM HCl was added. The reaction was initiated by adding peptide substrate Suc-Ala-Ala-Cis-Pro-Phe-pNA in LiCl/TFE solution with rapid inversion. The absorbance of p-nitroaniline was monitored at 390 nm until the reaction was complete (around 1 min). The final concentration of LiCl in the assay was 20 mM, and TFE was present at a concentration of 4% (v/v). Absorbance readings were collected every second using a spectrophotometer. For inhibition assessment, 5 ml of the tested compound in dimethyl sulfoxide (DMSO) was added to the cyclophilin solution in the assay buffer. Cyclosporine A (CsA, Sigma-Aldrich) was used as a positive control of PPIase inhibition in all measurements. The percentage inhibition of cyclophilin PPIase activity was calculated from the slopes, and the IC_50s_ values obtained represent the mean ± standard deviation of at least two independent measurements.

### SMCypI screening on FIPV infection

CRFK cells were plated in a 96-well plate at the density of 2.10^4^ cells/well and incubated for 24 hrs. Cells were infected for 1 hr with FIPV at MOI 1 in the presence of 50µM or 10µM of SMCypI or CsA respectively. Inoculum was removed after 1 hr and inhibitors at either 50μM for SMCypI or at 10μM for CsA were added for 24 supplemental hrs. The supernatants were collected and kept at -80°C prior to end-point dilution assays.

### FIPV titration by end-point dilution assay

CRFK cells were plated in a 96-well plate at the density of 2.10^4^ cells/well and incubated for 24 hrs. Dilutions of viral supernatants ranging from 10^-1^ to 10^-6^ in eight replicates per dilution were distributed in the CRFK plate. Viral inoculum was removed 2 hrs after infection and cells were incubated for 24 supplemental hrs at 37°C. Afterwards, cells were incubated for 10 minutes at room temperature in absolute ethanol, then in 70% ethanol, and FIPV antigens were stained by immunofluorescence using ascites liquids from an infected cat (dilution 1/500) followed by incubation with anti-cat A-488 secondary antibody (Jackson Immunoresearch). Wells containing fluorescent cells were counted. The 50% infectious dose (ID_50_) was calculated using the Spearman-Karber method. Results were the means of ≥ 2 experiments performed in triplicates.

### Cytotoxicity assay

Cells were plated in a 96-well plate at the density of 2.10^4^ cells/well and incubated for 24 hrs. Culture medium was replaced by serial dilutions of culture media containing three replicates of different concentrations (ranging from 25μM to 0.9μM) of F83233, F832 and F833 molecules, DMSO (Dimethyl Sulfoxide) or media alone as controls. Plates were incubated for 24 hrs at 37°C. Cytotoxicity was evaluated with the CellTiter-Glo Cell viability Assay kit (Promega).

### Dose-responses of SMCypI in CRFK, PK15 and Vero cells

CRFK, PK15 and Vero cells were plated in a 96-well plate at the density of 2.10^4^ cells/well and incubated 18 hrs at 37°C. Cells were infected with FIPV (MOI 1), TGEV (MOI 0.5) and PEDV (MOI 0.5) respectively. Viral inoculation was performed in the presence of F83233, F832 and F833 at concentrations ranging from 0.1μM to 25μM for FIPV assays and from 0.78µM to 12.5µM for TGEV and PEDV. Two hrs after viral inoculation, fresh medium was added with the same concentrations of inhibitors. Antiviral effect was measured after further 24 hrs for FIPV and PEDV infection and 48 hrs for TGEV. The experiments were repeated twice and each condition was tested in triplicates.

### Time-of-drug addition assay

CRFK cells were plated in a 96-well plate at the density of 2.10^4^ cells per well and incubated for 24 hrs. Cells were infected with MOI 1 of FIPV in the presence of 5µM F83233 for the 0 h post-infection (p.i.) condition or in presence of DMSO. After viral inoculation, culture medium was removed and fresh medium containing 5μM of F83233 was added at different times of the infection (1-, 3-, 6-, 9- or 12 hrs). Cells were incubated for supplemental 24 hrs at 37°C. Cell supernatants were kept at -80°C prior to end-point dilution assays to measure the effect of F83233 on FIPV infectivity in the different conditions. The experiment was repeated three times and each condition was tested in triplicates.

### Detection of TGEV and PEDV viral antigens by immunofluorescence

Fixed PK15 and Vero cells were processed similarly. TGEV infection was detected with a home-made anti-Spike antibody (called 51.13, dilution 1/10.000) from mouse and with an anti-mouse Alexa-Fluor 555 secondary antibody. PEDV infection was detected with polyclonal antibodies from pig (kind gift of Drs. Y. Blanchard and M. Contrant, Anses, Ploufragan, dilution 1/300) and with an anti-pig Alexa-Fluor 488 secondary antibody (SouthernBiotech). Active PEDV replication was revealed with a mouse antibody directed against dsRNA (J2 from Cell Signaling Technology, dilution 1/1.000) followed by incubation with an anti-mouse Alexa-Fluor 555 secondary antibody. Cells were counterstained with DAPI. Secondary antibodies (except the anti-pig) were used at a 1/800 dilution and are from Molecular Probes (Thermo Fischer Scientific).

### Quantification of TGEV and PEDV infected areas and nuclei/syncytium

For each concentration of F832, F833 and F83233, 3 to 5 pictures were taken with an epifluorescence microscope (Zeiss) with a 10X objective. The area of staining was measured with ImageJ. The number of nuclei/syncytia in PEDV-infected cells was manually counted using ImageJ.

### Alignment of amino acids sequences of cyclophilin A from different mammals

The cyclophilin sequences from human (P62937), pig (P62936), monkey (P62938) and cat (Q8HXS3) were recovered from UniProt (UniProt, http://www.uniprot.org/uniprot/), aligned with ClustalW and processed with ESPript 3.

### *In silico* modelling and docking

The search for ligand-cyclophilin 3D crystal complexes was performed using the @TOME-3 server^46^ (https://atome.cbs.cnrs.fr/ATOME_V3/index.html). Ligand files were generated with MarvinSketch 6.2.2 for SMILES and Grade server for mol2 (https://grade.globalphasing.org/cgi-bin/grade2_server.cgi). Docking simulation of F83233 in complex with pig cyclophilin A was performed using @TOME-3 server with an anchor of PDB 4J5C. The images were generated using PyMOL and MarvinSketch.

## Results

### Screening of non-peptidic small-molecule cyclophilin inhibitors (SMCypI) on Feline Infectious Peritonitis Virus (FIPV) infection

As cellular cyclophilins facilitate the replication of numerous viruses, we previously developed SMCypI to obtain broad-spectrum antiviral agents^45^. We have also previously shown that our first SMCypI series exhibited a potent antiviral activity on hepatitis C virus replication *in vitro*^47^, but that they were only modest to reduce the replication of the HCoV-229E CoV, with EC_50s_ ranging from 7 to 71µM^45^. SMCypI are non-peptidic compounds with a common backbone on which various chemical moieties can be added at three distinct sites referred as R1, R2 and R3 (**Fig. 1A**), thus allowing to routinely generate new molecules (**Table 1**). We thus aimed to find more potent anti-CoV compounds in our enriched library.

**Table 1:**
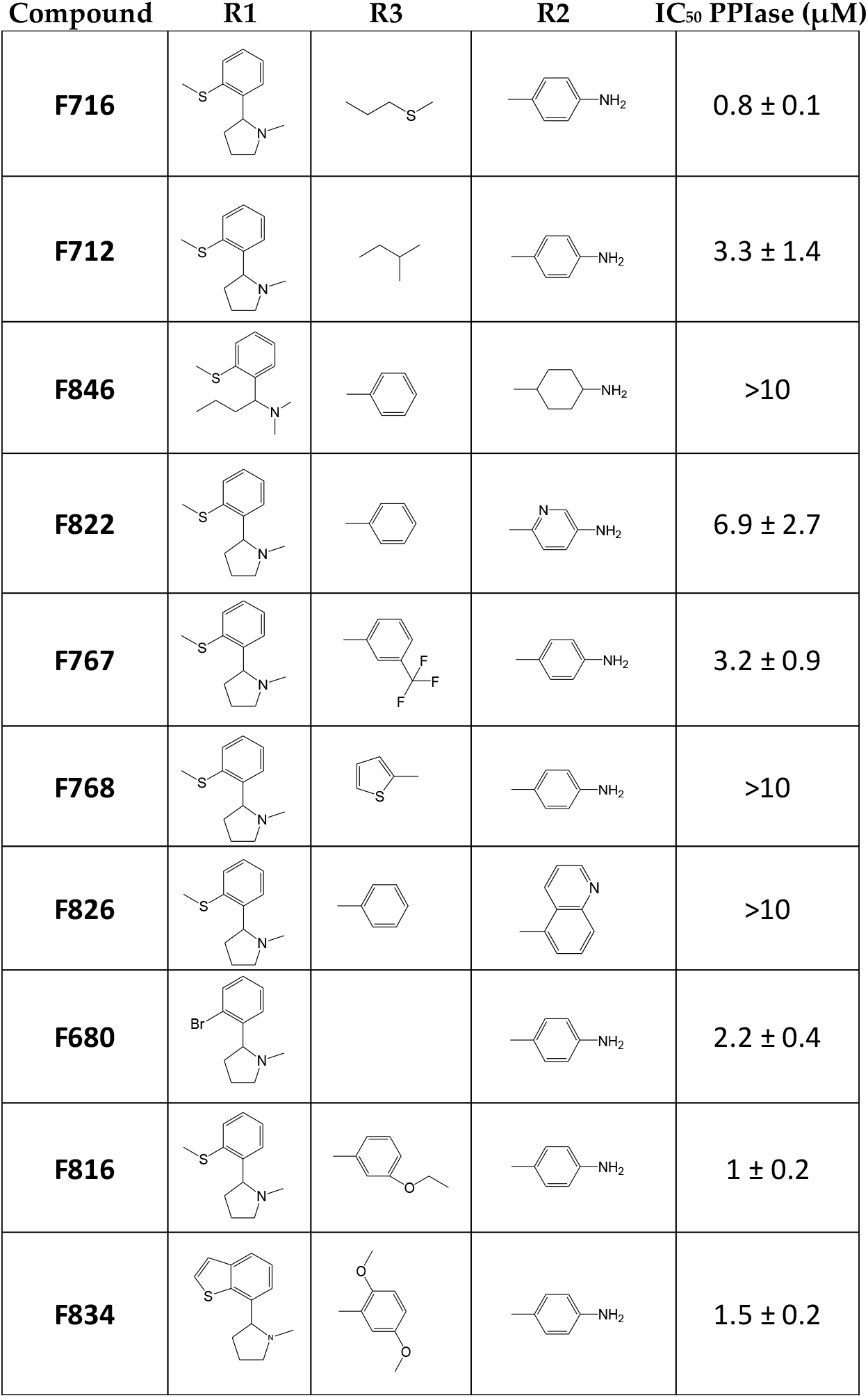
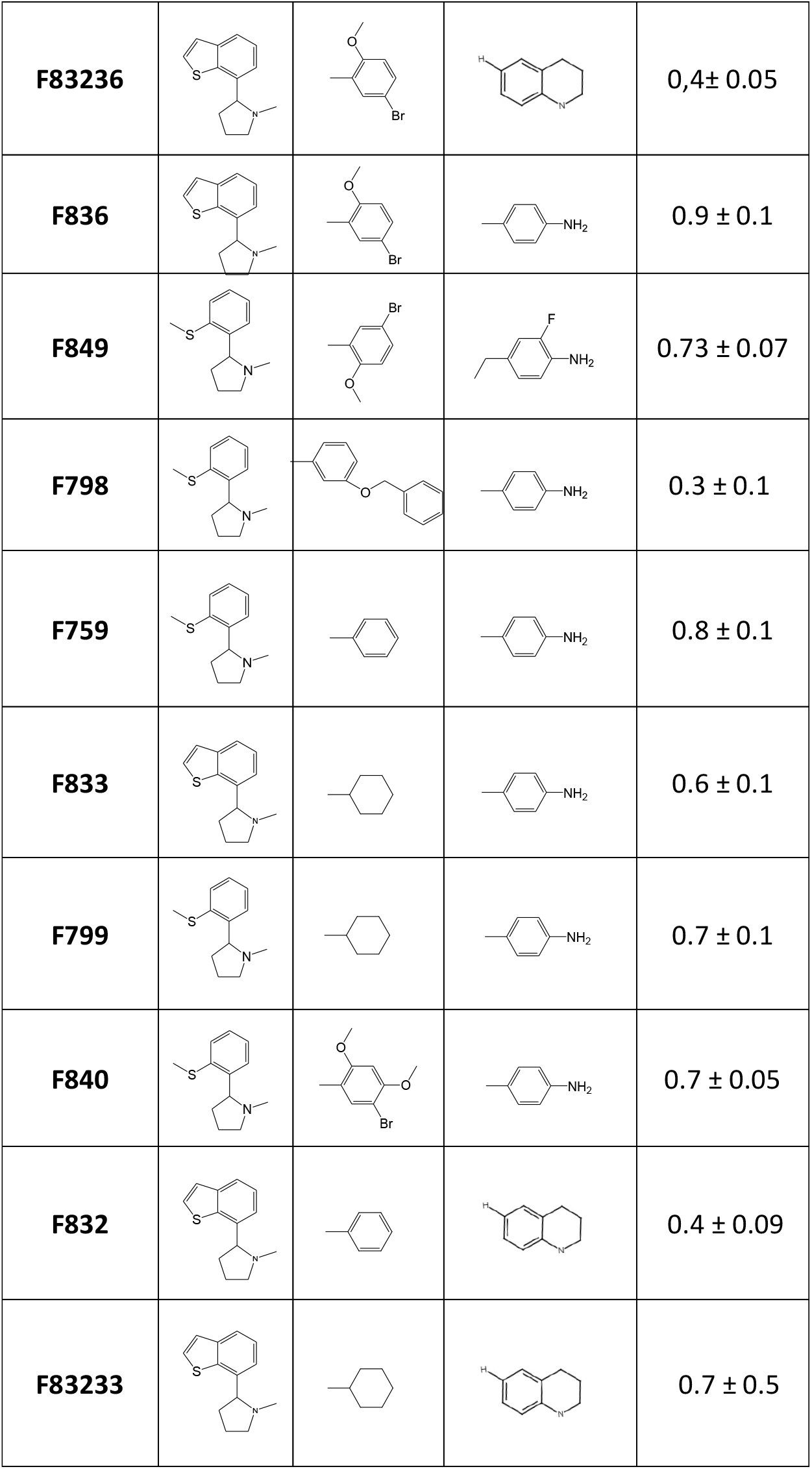
Structures and anti-PPIase activities of SMCypI.

**Figure 1:**
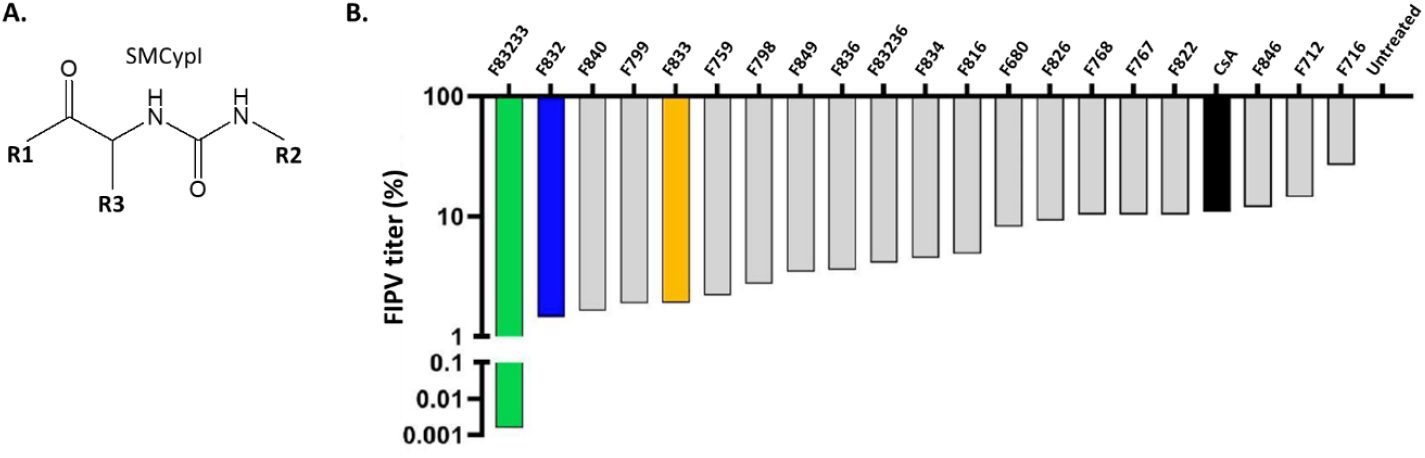
Screening of non-peptidic small-molecule cyclophilin inhibitors (SMCypI) on Feline Infectious Peritonitis Virus (FIPV) infection. **A**. SMCypI are composed of a common backbone with various substitutions that could be added at the R1, R2 and R3 regions, allowing to generate a library with an important number of compounds. **B**. SMCypI are more effective than CsA. FIPV viral titer was measured 24 hrs after treatment with 50µM of 20 different SMCypI. Cyclosporin A (CsA) (in black) was used at the final concentration of 10µM. Results are normalized with untreated infected cells.

We performed a screen of 20 different SMCypI (**Fig. 1B** and **Table 1**) for their ability to inhibit Feline Infectious Peritonitis Virus infection (FIPV, 79-1146 strain) in feline CRFK cells at the final concentration of 50µM. We also used 10µM of cyclosporine A (CsA, black bar), a peptidic cyclophilin inhibitor with a reported anti-CoV activity, including FIPV^38^. We observed that all the tested molecules modestly impacted the infectious titers of FIPV, except the F83233 compound (green bar) that inhibited FIPV titer by more than 4 Log_10_ (**Fig. 1B**).

### The increased anti-FIPV effect of F83233 compared to F832 and F833 is unrelated to cyclophilin A enzymatic activity inhibition

SMCypI F83233 exhibits chemical moieties from F832 and F833 (blue and orange bars on **Fig. 1B**, respectively) at R2 and R3 positions, respectively (**Fig. 2A** and **Table 1**). We first confirmed the results from our screen by measuring the virus titer in cell supernatants after FIPV infection in the presence of 25µM SMCypI. While F832 and F833 only modestly inhibited FIPV infectivity by ≈0.5 Log_10_ (**Fig. 2B**), F83233 decreased by ≈3 Log_10_ the virus titer at this concentration. We measured the ability of these three compounds to block the enzymatic activity of human cyclophilin A in an *in vitro* assay to assess if this increased antiviral potency of F83233 was linked to an improved blockade of cyclophilin function (**Table 1**). However, we observed no significant differences with IC_50s_ of 0.7 ± 0.5µM for F83233, 0.4 ± 0.09µM for F832 and 0.6 ± 0.1µM for F833 (**Table 1**). As apparent antiviral effect could sometimes be due to cellular toxicity, we treated CRFK cells for 24 hrs with increasing concentrations of the three compounds, and observed a low cytotoxicity of the drugs up to 25µM, that was not different between F83233 and F832/F833 (**Fig. 2C**), an observation that was similar in the simian Vero cell line (Softic *et al*. submitted). Overall, the increased activity of F83233 compared to its “ parental” molecules is not due to an ameliorated anti-PPIase activity on cyclophilin A or to cellular toxicity in our experimental conditions.

**Figure 2:**
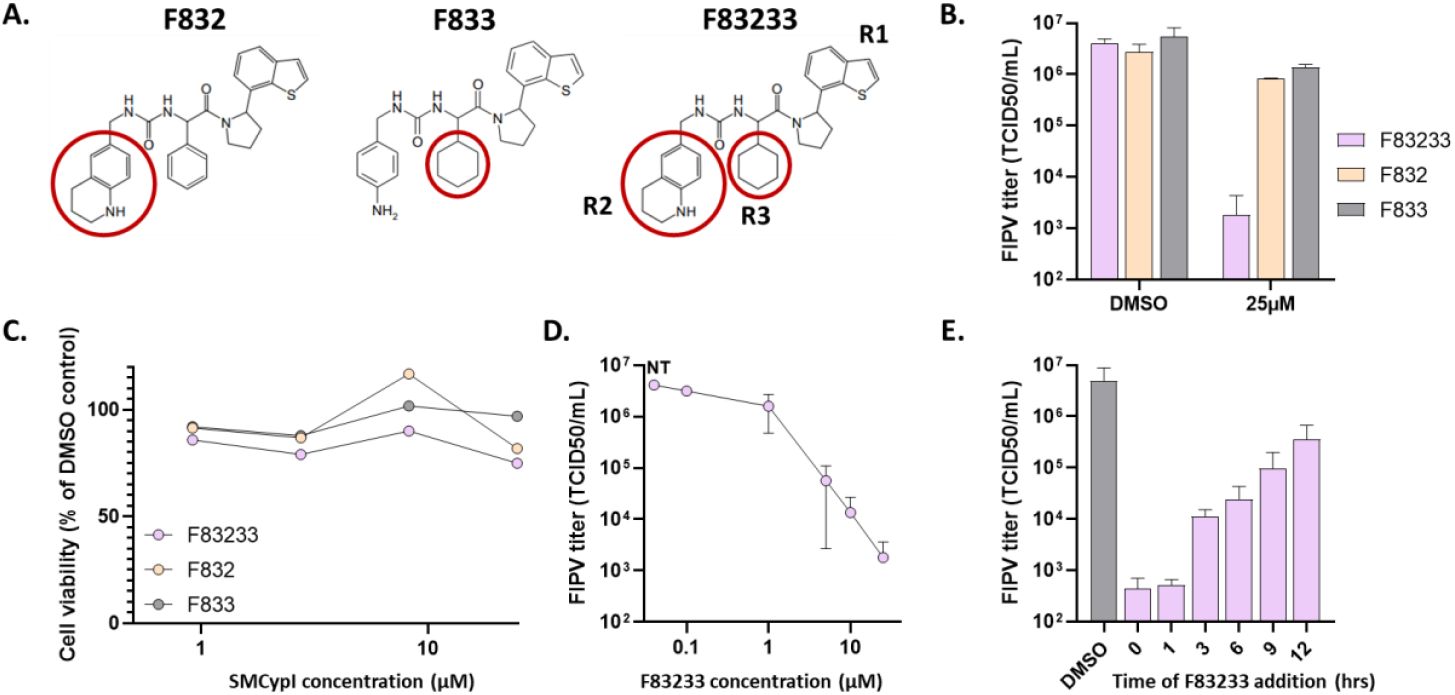
Characterization of the anti-FIPV activity of SMCypI F83233. **A**. Chemical structures of F832, F833 and F83233 molecules. F83233 harbours, at R2 and R3 positions, chemical moieties from F832 and F833 respectively (red circles). **B**. Antiviral effect of 25µM of F83233 compared with those of F832 and F833 molecules was assessed by measuring FIPV viral titer in cell supernatants by TCID_50_ after treatment during 24 hrs with the drugs or DMSO. n = 2 independent experiments. **C**. Cytotoxicity assays. Measure of ATP release after treatment with increasing concentrations of F83233, F832 and F833. **D**. Dose-responses. FIPV viral titers were measured in cell supernatants after treatment with escalating concentrations of F83233. n = 2 independent experiments. **E**. Time of addition assays. F83233 (5µM) was added to the cells at various times post-infection (between 0 and 12 hrs) and FIPV titers from cell supernatants were measured in the different conditions. n = 3 independent experiments. N.T: Non-Treated.

### Characterization of the anti-FIPV activity of F83233

To characterize the anti-FIPV activity of F83233, we first quantified FIPV infectious titers in the presence of escalating doses of this SMCypI (**Fig. 2D**). We demonstrated a strong inhibition of FIPV infectivity with a reduction in viral titer by approximately 2 Log_10_ at 5 µM. Furthermore, the maximal inhibitory effect of F83233 was observed as a decrease of approximately 3-4 Log_10_ at 12.5 µM. Then, to better understand its mechanism of action on the FIPV replication cycle, we added 5µM of F83233 at different times post-infection (**Fig. 2E**). The maximal antiviral effect of F83233 was observed when the molecule is added simultaneously with the virus or 1 hr later. Its antiviral efficiency decreased by more than 1 Log_10_ between 1 hr and 3 hrs post-infection. From 3 hrs to 12 hrs post-infection, F83233 continuously lost antiviral efficiency but was still effective 12 hrs post-infection with ≈1 Log_10_ of inhibition as compared to the DMSO control. Altogether these data indicate that F83233 mainly blocked an early step of the FIPV replication cycle.

#### F83233 inhibits infection by Transmissible Gastroenteritis Virus (TGEV) in porcine cells

We next evaluated if F83233 could also inhibit infections by other α-CoVs of veterinary interest. We first studied infection by porcine Transmissible Gastroenteritis Virus (TGEV, Purdue strain) in porcine kidney PK15 cells (**Fig. 3A** and **3B**). As for FIPV, we compared the antiviral potency between F83233 and F832/F833. While F832 and F833 had no significant effect on TGEV infection even at 12.5µM, F83233 was active at 1.5µM and was associated with an almost complete disappearance of infected cells at 6.25µM and 12.5µM (**Fig. 3A** and **3B**).

**Figure 3:**
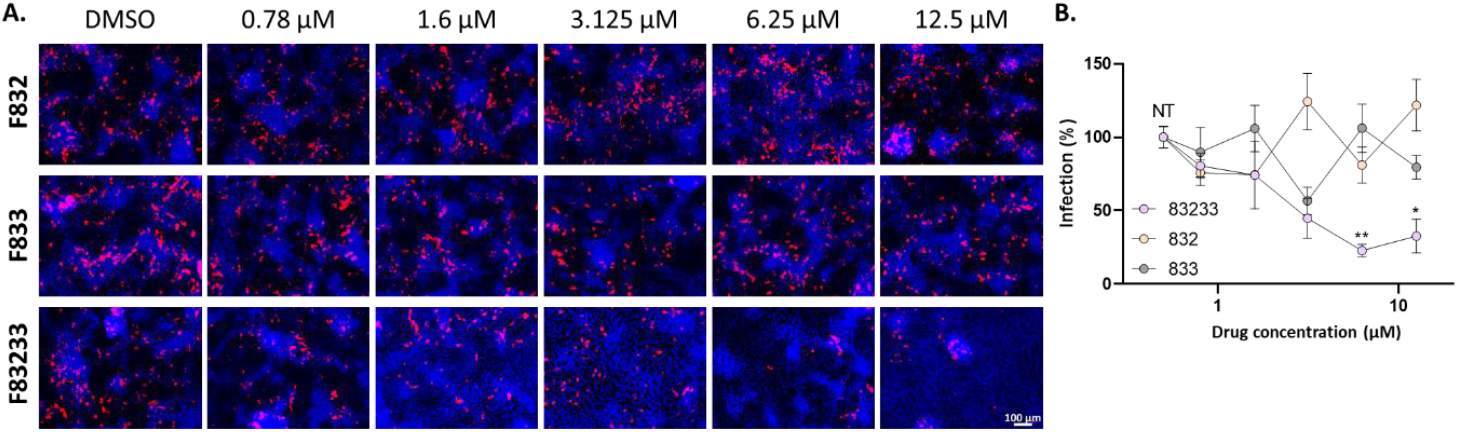
Antiviral activity of SMCypI against porcine Transmissible Gastroenteritis Virus (TGEV) in porcine cells. **A**. PK15 (porcine kidney) cells were infected by TGEV (Purdue strain) at a MOI of 0.5 for 48 hrs in presence of increasing concentrations of SMCypI or only DMSO. Viral infection was detected by immunofluorescence using an anti-Spike monoclonal antibody and an anti-mouse A-555 secondary antibody (red). Cell nuclei were counterstained with DAPI. Objective: 10X. **B**. Quantification of TGEV inhibition by SMCypI. A total of 3-5 images was captured for each condition, with two images per well. The area of infection was then quantified using ImageJ. Figure from one experiment representative of n = 2 independent experiments with 2 independent wells. N.T.: Non-Treated. *: p<0.05, **: p<0.01 (One-Way Anova followed by Kruskal-Wallis test).

#### F83233 inhibits infection by Porcine Epidemic Diarrhoea Virus (PEDV) in simian cells

To study the broad-spectrum potential of F83233, we tested its antiviral effect on the replication of Porcine Epidemic Diarrhoea Virus (PEDV, CV777 strain), which is more genetically distant from FIPV. In simian Vero cells infected by PEDV, we observed a modest antiviral potency of F832 and F833, while F83233 allowed to observe an important reduction of infection at 3µM, and a maximal antiviral effect between 3 and 6µM (**Fig. 4A** and **4B**).

**Figure 4:**
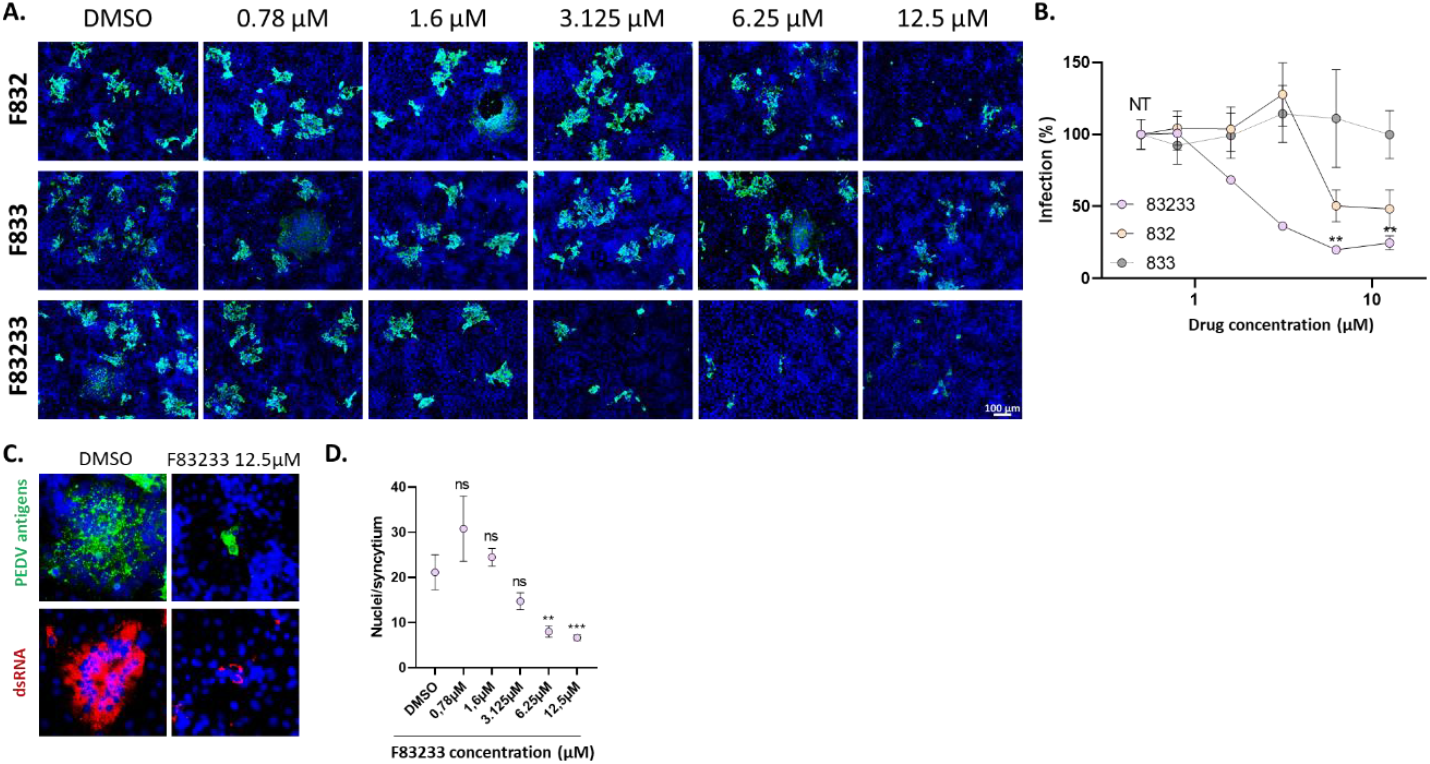
Antiviral activity of SMCypI against Porcine Epidemic Diarrhoea Virus (PEDV) in simian cells. **A**. Vero (monkey kidney) cells were infected with PEDV (CV777 strain) at an MOI of 0.5 for 24 hrs in presence of increasing concentrations of SMCypI or only DMSO. Viral infection was detected by immunofluorescence using anti-PEDV porcine polyclonal antibodies and an anti-pig secondary antibody (green). Cell nuclei were counterstained with DAPI. Objective: 10X. **B**. Quantification of PEDV inhibition by SMCypI. A total of 3-5 images were captured for each condition, with two images per well. The area of infection was then quantified using ImageJ. Figure from one experiment representative of n = 2 independent experiments with 2 independent wells. N.T.: Non-Treated. **: p<0.01 (One-Way Anova followed by Kruskal-Wallis test). NT: Non-Treated. **C**. SMCypI treatment reduced drastically syncytia formation. Infected Vero cells were treated with 12.5µM of F83233 and stained after fixation with anti-PEDV polyclonal antibodies and an anti-pig secondary antibody (green), or with a mouse antibody that recognizes dsRNA structures and an anti-mouse secondary antibody (red). Objective: 10X. **D**. Quantification of the number of nuclei per syncytia after treatment with increasing doses of F83233. The number of nuclei per syncytium was counted in four different fields of cells (2 pictures per well). Figure from one experiment representative of n = 2 independent experiments with 2 independent wells. ns: not significant. ***: p<0.001 (One-Way Anova followed by Kruskal-Wallis test).

In cultured cells, a number of CoVs are characterized by an atypical cytopathic effect, *i*.*e*. the formation of multinucleated cells referred as syncytia, as a result of a massive cell-cell fusion triggered by the viral Spike protein^48–50^. In Vero cells, PEDV appears to be one of the most fusogenic CoV and we observed in our experimental conditions the presence of several syncytia containing up to 100 nuclei (**Fig. 4C**, DMSO condition). Using an antibody that recognizes dsRNA structures (that are formed during the replication of viruses with RNA genomes of positive polarity), we observed that syncytia are an active site of RNA replication, with dsRNA spots located around the packed nuclei (**Fig. 4C**).

Using 12.5µM F83233, we noticed that the SMCypI drastically decreased the size of the PEDV-induced syncytia and the number of nuclei/syncytium (**Fig. 4C** and **4D**), and markedly impacted the dsRNA staining corresponding to viral genome replication in the syncytia (**Fig. 4C**).

## 4. Discussion

The regular emergence of new coronaviruses (CoVs) in humans and animals necessitates the urgent development of antiviral tools with a broad pan-CoV spectrum. Two main antiviral methodologies are commonly employed: either to target specifically viral proteins or to target cellular proteins that are necessary for viral replication. Whilst the initial approach is more frequently used, it is not without its drawbacks. Firstly, it is frequently virus-specific, and secondly, it is difficult to extrapolate its antiviral efficiency to other CoVs. Finally, it gives rise to the concern of the emergence of antiviral-resistant mutants. Recently, a nucleoside analogue, GS141524, a metabolite of the antiviral prodrug remdesivir, has been employed to treat FIP disease, with considerable success^8,9^. However, this strategy is not efficacious against SARS-CoV-2 for example^51^. In this study, we chose to target the cellular cyclophilin A (encoded by PPIA gene), the most abundant cyclophilin. As demonstrated in previous studies, this protein has been shown to be necessary for the replication of multiple CoVs from different genera^33^. Furthermore, evidence has demonstrated that cyclosporine A (CsA), a macrocyclic peptide composed of 11 amino acids, forms a complex with cyclophilin A, thereby exhibiting an antiviral effect against FIPV, TGEV and PEDV at least^38,42,52^. Nevertheless, the immunosuppressive effect of this compound prevents its utilisation as an antiviral therapeutic agent.

We previously developed low molecular weight, non-immunosuppressive cyclophilin-inhibiting molecules (SMCypI). Subsequently, the anti-HCoV-229E activity of SMCypI was characterised, exhibiting a modest antiviral effect^45^. However, the relatively simple chemistry of these molecules allows to envisage numerous substitutions on their common backbone to generate a more diverse library (**Fig. 1A** and **Table 1**). With this approach, we described a new antiviral molecule, F83233, with a more efficient antiviral activity (around 3 Log_10_ viral reduction at 10*μ*M) than CsA against FIPV (**Fig 1B** and **2D**). The potent antiviral activity of F83233 results directly from the easy possibility to “ mix” groups of two “ parental” compounds such as F832 and F833. These data pave the way for the screening of more SMCypI, and could lead to structure-activity relationship studies for the design of molecules even more effective than F83233 against CoVs (as well as other viruses) multiplication.

For some viruses whose replication is cyclophilin-dependent, it has been demonstrated that the PPIase enzymatic activity is necessary^33^. For instance, the genome replication of the hepatitis C virus (HCV) absolutely requires the PPIase activity of cyclophilin A^33,53^. Similarly, we observed a very good correlation between the ability of SMCypI to inhibit the PPIase activity of human cyclophilin A and their anti-HCV potency^47^. In our present study, SMCypI with little or no ability to block cyclophilin A PPIase function (F846, F768, F826) also showed a modest anti-FIPV effect (**Fig. 1B** and **Table 1**). By contrast, compounds with a more pronounced antiviral potency (F832, F840, F799, F833, F759) blocked cyclophilin A PPIase activity with sub-micromolar IC_50s_. However, the massive increase in anti-CoV activity of F83233 compared to these molecules is not linked to an improved potency on the PPIase activity of cyclophilin A, as F83233 exhibits a similar IC_50_ (**Table 1**). In the same line, CsA at 10µM was less effective than F83233, whereas its IC_50_ on the PPIase activity is 2 Log_10_ lower than F83233. Thus, the antiviral effect of SMCypI on CoVs multiplication may be more complex than that observed on HCV, and these molecules could be very useful to decipher the interactions between cyclophilins and CoVs.

The development of broad-spectrum molecules can help to prevent future pandemic^54^. While specific antivirals or vaccines are being developed, this can for example provide a first line of defence in the event of viral emergence. Antivirals targeting host factors may thus act on a wide range of viruses and animal species thanks to their conservation. In this study, the F83233 demonstrated robust antiviral efficacy against both FIPV and TGEV, which are genetically related. In addition, it was found to inhibit PEDV, a distinct α-CoV that exhibited only 31% and 29 % amino-acid homologies of the Spike and N proteins respectively, when compared with TGEV. The broad activity of F83233 on PEDV, FIPV and TGEV in simian, feline and porcine cells, respectively, demonstrates that SMCypI are serious leads for considering treatments that can be generalized to several animal species, and ultimately prevent the zoonotic risk posed by a mammalian CoV. This broad activity is consistent with the very strong conservation of cyclophilin A and its active PPIase site across mammals (**Fig. 5A**). The 3D reconstruction of the complex made by F83233 and cyclophilin A shows identical interactions with cyclophilin A accross the different species (human, porcine [shown on **Fig. 5B**]), simian and feline). This paves the way for the study of SMCypI antiviral activity on animal CoVs infecting other species.

**Figure 5:**
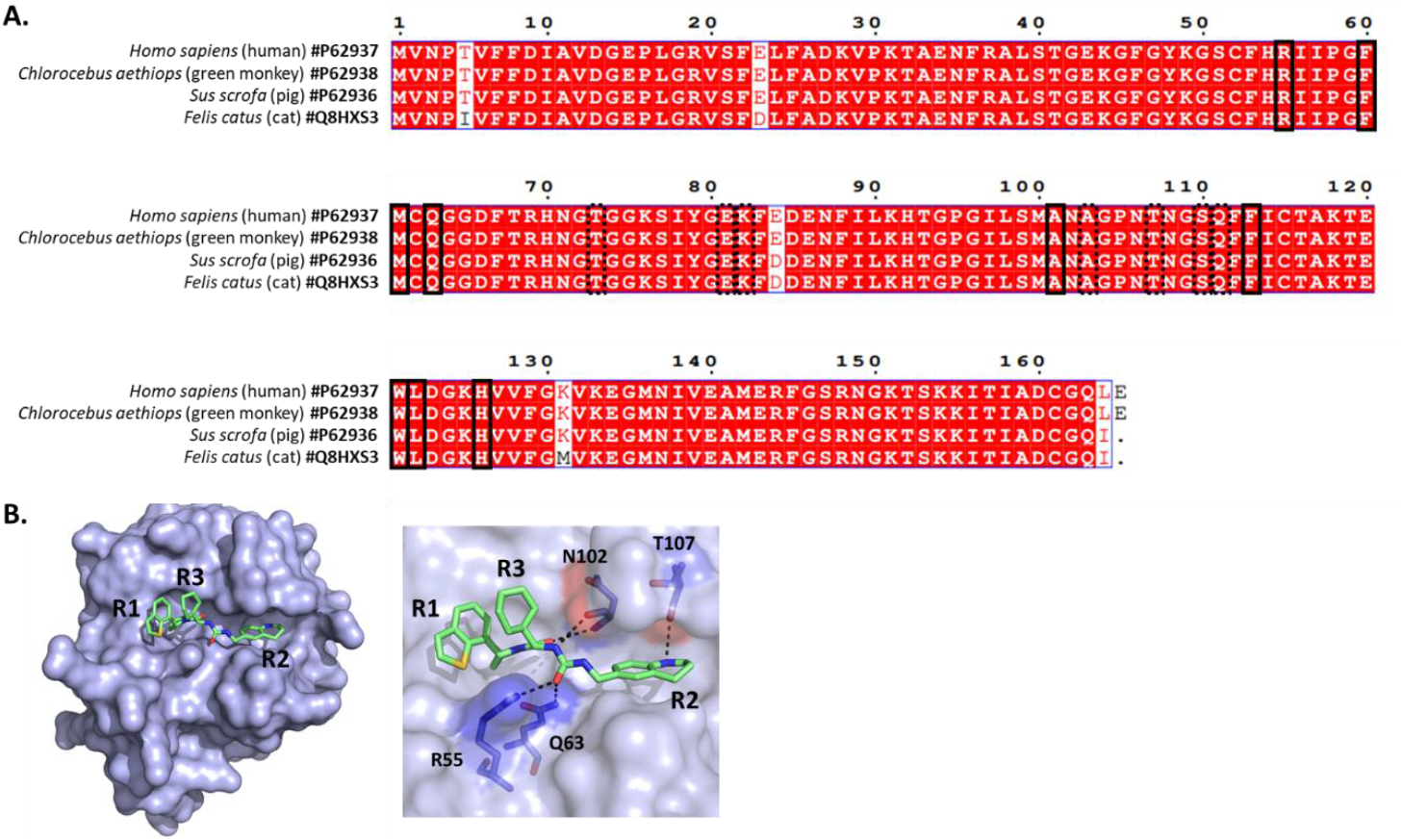
High-degree of conservation of cyclophilin A across mammals. **A**. Sequences of cyclophilin A (PPIA gene) were obtained from Uniprot, aligned with ClustalW and processed with ESPript 3. The alignment shows high-degree conservation of cyclophilin A from human, monkey, pig and cat. Amino-acids of the PPIase active site are in dashed rectangles while amino-acids that define the S2 “ gatekeeper” pocket (that regulated the substrate specificity^31^) in dotted rectangles. **B**. In silico modelling and docking. The cyclophilin A sequences from human (P62937), pig (P62936), monkey (P62938) and cat (Q8HXS3) were recovered from UniProt. The ligand (F83233, in green)-cyclophilin 3D crystal complex was performed using the @TOME-3 server^46^ (https://atome.cbs.cnrs.fr/ATOME_V3/index.html). Ligand files were generated with MarvinSketch 6.2.2 for SMILES and Grade server for mol2. Docking simulation was performed using @TOME-3 server with an anchor of PDB 4J5C. The images were generated using PyMOL and MarvinSketch. The pig cyclophilin A is shown on the figure.

## Acknowledgments

This work was supported by Anses grant «CycloCoV». MD was supported by DIM1 Health (Region Ile de France). MK is supported by a grant from Anses. QN obtained grants from INRAE Animal Health Department, Université Paris-Saclay and LSH graduate school “ OI Microbes” , and is supported by the Paris-Saclay ANR PIA funding: ANR-20-IDEES-0002 grant. We are grateful to Dr Nicolas Meunier for critical reading of the manuscript.

## Authors contributions

Conceptualization: SLP, QN, AAB. Investigation: MD, MK, HH, LS, LB, JFG, AAB, QN and SLP. Writing: MD, AAB, QN and SLP, with input from all authors.

